# Activation of Group II Metabotropic Glutamate Receptors in the Basolateral Amygdala Inhibits Reward Seeking Triggered by Discriminative Stimuli

**DOI:** 10.1101/2024.10.11.617854

**Authors:** Mandy Rita LeCocq, Amélie Mainville-Berthiaume, Isabel Laplante, Anne-Noël Samaha

**Affiliations:** Department of Pharmacology and Physiology, Faculty of Medicine, Université de Montréal, Montreal, QC, Canada; Neural Signaling and Circuitry Research Group (SNC), Faculty of Medicine, Université de Montréal, Montréal, QC, Canada; Center for Biomedical Innovation (CIB), Université de Montréal, Montréal, QC, Canada; Center for Interdisciplinary Research on the Brain and Learning (CIRCA), Université de Montréal, Montréal, QC, Canada; Center for Studies in Behavioural Neurobiology, Montréal, QC, Canada

**Keywords:** Conditioned stimulus, Appetitive, Reinforcement, Pavlovian conditioning, Conditioned approach, Cue

## Abstract

Reward-associated cues are essential in guiding reward-seeking behaviours. These cues include conditioned stimuli (CSs) which occur following seeking actions and indicate reward delivery, and discriminative stimuli (DSs) which occur response-independently and signal reward availability. Metabotropic group II glutamate (mGlu_2/3_) receptors in the basolateral amygdala (BLA) modulate CS-guided reward seeking; however, their role in DS effects is unknown. We first developed a procedure to assess DS and CS effects on reward seeking in the same subjects within the same test session. Female and male rats first self-administered sucrose during sessions where discriminative stimuli signaled periods of sucrose availability (DS+) and unavailability (DS-). During DS+ presentations, active lever presses produced sucrose paired with a CS+. During DS-presentations, active lever presses produced a CS- and no sucrose. Across 14 sessions, rats learned to load up on sucrose during DS+ presentation and inhibit responding during DS-presentation. We then compared the effects of intra-BLA microinfusions of the mGlu_2/3_ receptor agonist LY379268 on cue-evoked sucrose seeking during an extinction test (no sucrose) where the DSs and CSs were presented response-independently. Before test, rats received intra-BLA microinjections of artificial cerebrospinal fluid (aCSF) or LY379268. Under aCSF, only the DS+ and DS+CS+ combined triggered increases in reward-seeking behaviour. The CS+ alone was ineffective. Intra-BLA LY379268 reduced sucrose seeking triggered by the DS+ and DS+CS+ combination. Thus, using a new procedure to test reward seeking induced by DSs and CSs, we show that BLA mGlu_2/3_ receptor activity mediates the conditioned incentive motivational effects of reward predictive DSs.

## Introduction

An essential feature of animal behaviour is the capacity to respond flexibly to environmental stimuli (e.g., smell) that predict rewards required for survival (e.g., food). Such reward cues can guide adaptive behaviours like seeking out food; however, they can also promote maladaptive reward seeking that contributes to eating disorders, addiction, and pathological gambling [1–5].

When animals are seeking rewards, distinct environmental cues hold different associative meanings. Conditioned stimuli (CSs) occur as a consequence of reward-seeking actions and are paired with reward delivery [6,7] (e.g., the sensation of water touching a thirsty animal’s mouth as it drinks). In comparison, discriminative stimuli (DSs) signal reward availability, independent of any reward-seeking actions [6,8] (e.g., unexpectedly hearing the sound of a water stream). Thus, DSs are unique in that they are often encountered accidentally, before an individual is even aware of desiring any particular reward, and thus are at play before, during, and after reward-seeking actions.

When presented response-contingently, both CSs and DSs can act as conditioned reinforcers of instrumental responding [9–15]. However, when presented response-independently, a CS is rather ineffective in triggering increases in reward seeking [9,15–19], whereas a DS is very effective in doing so [15,16,20–23] — even following extended abstinence and repeated testing [20,22,24]. Despite the unique properties of DSs, comparatively little is known about the neurobiological underpinnings of DS versus CS effects on reward-seeking behaviour.

The basolateral amygdala (BLA) mediates the ability of reward-associated cues to guide reward seeking. Neuronal activity in the BLA and its projections contributes to both CS [10,25–31] and DS [16,32–34] effects on reward-seeking behaviour. The BLA contains a high density of mGlu_2/3_ receptors [35–37]. Located mainly extrasynaptically on presynaptic terminals, mGlu_2/3_ receptors suppress glutamate release into the synaptic cleft and reduce the excitability of projection neurons [38–40]. Glutamate neurotransmission in the BLA contributes to the ability of CSs to energize and reinforce reward seeking [41–44], and intra-BLA injection of the mGlu_2/3_ receptor agonist LY379268 reduces CS-triggered increases in reward seeking, presumably by reducing glutamate neurotransmission [45]. However, the extent to which mGlu_2/3_ receptors in the BLA contribute to DS effects on reward-seeking behaviour remains unknown.

Here, we compared the influence of intra-BLA infusions of the selective mGlu_2/3_ receptor agonist LY379268 on DS and CS effects on sucrose seeking. To this end, we developed a procedure to compare CS and DS effects in the same subjects within the same session, to identify unique and similar underlying neurobiological mechanisms.

## Methodology

### Subjects

Female and male Sprague-Dawley rats (n = 18 females 9 weeks old, n = 16 males 7 weeks old. Charles River, Raleigh – RMS, Barrier; R04) weighing 225 – 250 g on arrival were single-housed. After an initial 72 h, rats were handled daily by the experimenters. Cages contained sani-chip bedding and a nylabone (Bio-Serv, K3580) and unrestricted access to food (Charles River Laboratories, Saint-Constant, Quebec, Canada). Access to water was restricted when specified. Cages were held in a climate-controlled colony room (22 ± 1°C, 30 ± 10% humidity), on a reverse 12 h light/dark cycle (0830 h lights off). All experiments were conducted during the dark phase. All procedures followed the Canadian Council on Animal Care guidelines and were approved by the Université de Montréal. The timeline at the top of each figure illustrates the sequence of experimental steps. See Supplementary document for information on apparatus, stereotaxic surgery, sucrose habituation, magazine training, histology, and brain microinfusions.

### Autoshaping

Rats with a sign-tracking phenotype may respond more to reward-associated CSs, while rats with a goal-tracking phenotype may respond more to reward-predictive DSs [46–49]. As such, rats first received 6 autoshaping sessions (30-min/session) to identify their Pavlovian conditioned approach phenotype (i.e., sign-tracker, goal-tracker, intermediate). Sessions began with a lever extending for 8 s (the conditioned stimulus, CS), followed by the delivery of 0.1 ml of sucrose solution. Lever-sucrose pairings occurred on a VI 30-s schedule (possible intervals: 4, 9, 14, 19, 24, 29, 34, 39, 44, 49, 54 s) with 30 pairings/session. The lever CS was counterbalanced across the right- and left-side levers. This lever was the active lever used in following experimental phases. The behavioural procedure is illustrated in Figure 1A.

**Figure 1.**
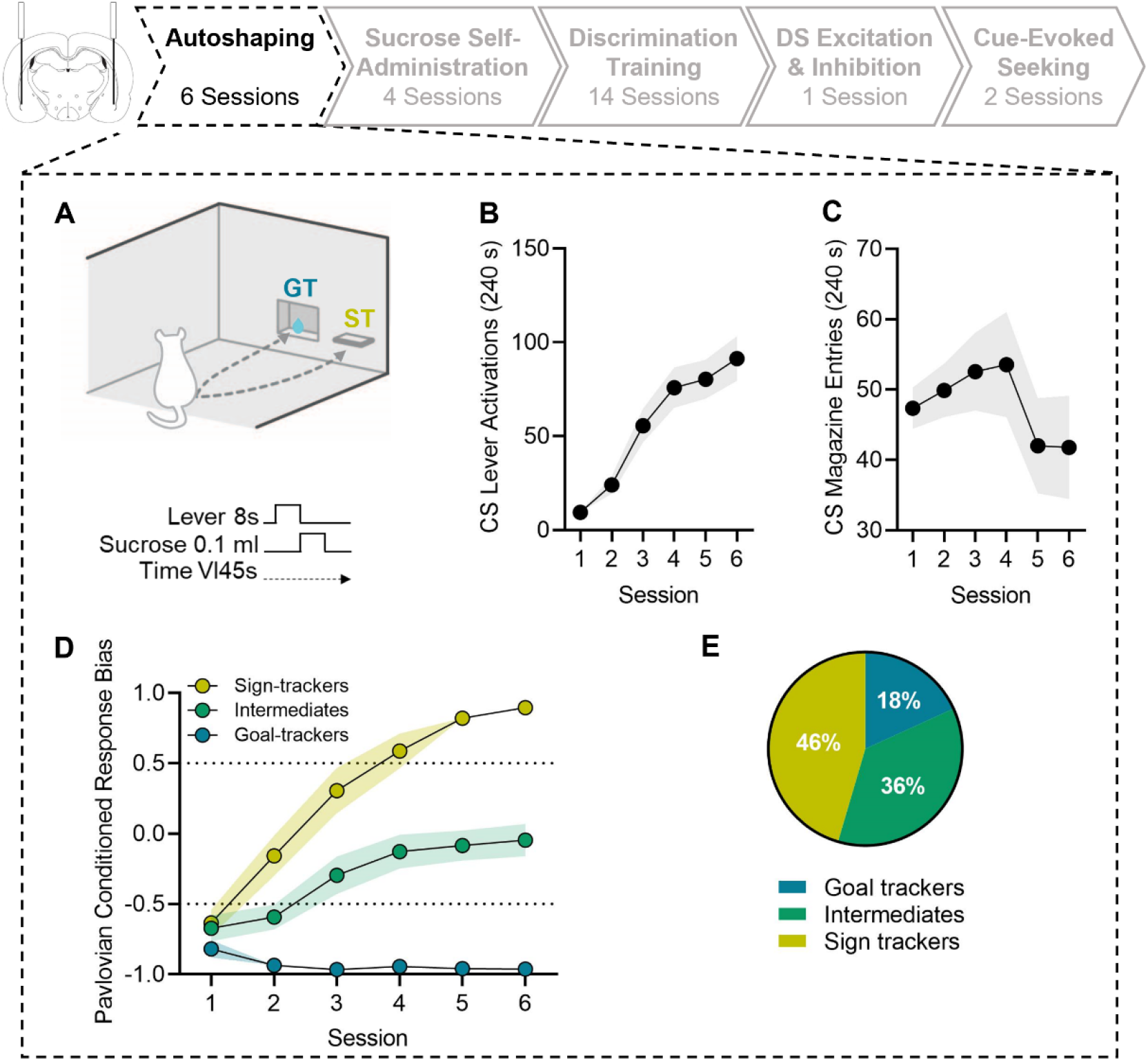
Acquisition of Pavlovian conditioned responding. **(A)** Schematic of the Pavlovian autoshaping procedure, adapted from Servonnet et al., 2023, with permission from Elsevier. Average (± SEM) **(B)** lever activations and **(C)** magazine entries made during the 8-s lever CS presentations. **(D)** Average (± SEM) response bias for sign-tracking (yellow circles), goal-tracking (blue circles), and intermediate (green circles) phenotypes. Dashed lines indicate the response bias score cut-offs (+0.50 and -0.50). **(E)** Percentage of phenotypes. *n* = 16 males, 17 females.

A response bias score was calculated to identify the rats’ Pavlovian conditioned approach phenotype, using data averaged over the last two autoshaping sessions [(CS lever presses - CS port entries) / (CS lever presses + CS port entries)]. A score of +0.50 or higher indicates sign-tracking, a score of -0.50 or lower indicates goal-tracking, and a score of -0.49 to +0.49 indicates intermediate responding [23,50].

### Sucrose Self-Administration

Following autoshaping, rats were trained to press a lever to self-administer sucrose in daily 1-h sessions. No cues were presented during this phase of training. Each session began with both levers extending. Pressing the active lever produced 0.1 ml sucrose and pressing the inactive lever had no programmed consequence (see Figure 2A).

**Figure 2.**
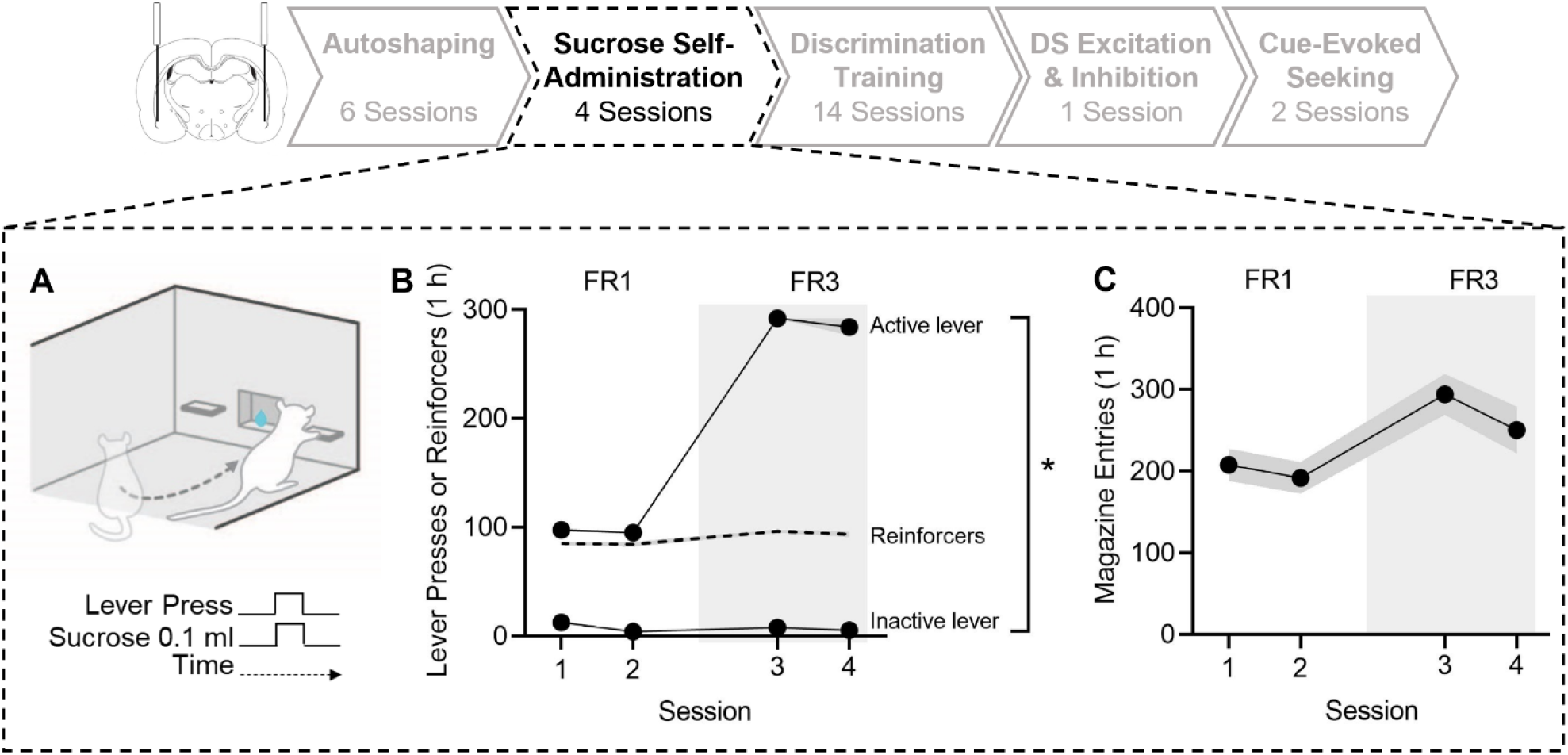
Acquisition of sucrose self-administration. **(A)** Schematic of the sucrose self-administration procedure, adapted from Servonnet et al., 2023, with permission from Elsevier. Average (± SEM) **(B)** active lever presses, inactive lever presses, reinforcers earned, and **(C)** magazine entries made, during FR1 and FR3 sessions. FR, Fixed ratio schedule of sucrose reinforcement. *Active lever presses > inactive lever presses across sessions. n = 16 males, 17 females.

The first sessions were conducted under a fixed ratio one (FR1) schedule of reinforcement. When rats earned 20 reinforcers and made twice as many active versus inactive lever presses for two consecutive sessions, they were moved to a FR3 schedule. When rats earned 30 reinforcers and made twice as many active versus inactive lever presses, they were moved to discrimination training. All rats met both sets of acquisition criteria after two sessions.

### Discrimination Training

Rats received 14 sessions (1 h/session) to learn to discriminate between a DS+ that signalled sucrose availability, and a DS- that signalled sucrose unavailability. They also learned that each sucrose delivery was paired with a CS+, while a CS- was never paired with sucrose. Table 1 describes the stimulus modality of each cue type. Stimulus modalities for the CS+ and CS- were counterbalanced, and the DS+ and CS+ were not explicitly above the active or inactive levers, as the active lever was counterbalanced across the right- and left-side levers.

**Table 1.** Cue modalities.

| Cue | Sample size | Stimulus type | Duration |
| --- | --- | --- | --- |
| DS+ | 34 | Cue light right side, front wall | 1-min |
| DS- | 34 | Cue light top-center, back wall | 1-min |
| CS+ | 17 | Cue light left side, front wall + tone, back wall | 5-s |
|  | 17 | Cue light bottom-left, back wall + clicker, back wall | 5-s |
| CS- | 17 | Cue light left side of front wall, + tone back wall | 5-s |
|  | 17 | Cue light bottom-left, back wall + clicker, back wall | 5-s |

Sessions began with the active and inactive levers extending, and a 1-min DS+ presentation during which active lever pressing produced 0.1 ml sucrose paired with a 5-s CS+ under FR3. DS+ presentations were followed by a 1-min DS-presentation, during which active lever presses produced a 5-s CS-under FR3, but no sucrose delivery. This alternating DS presentation cycled without interruption until each cue was presented 30 times (Figure 3A). Five minutes before the 14th discrimination training session, rats received an intra-BLA microinjection of aCSF (0.5 ul/hemisphere) to habituate them to microinfusions.

**Figure 3.**
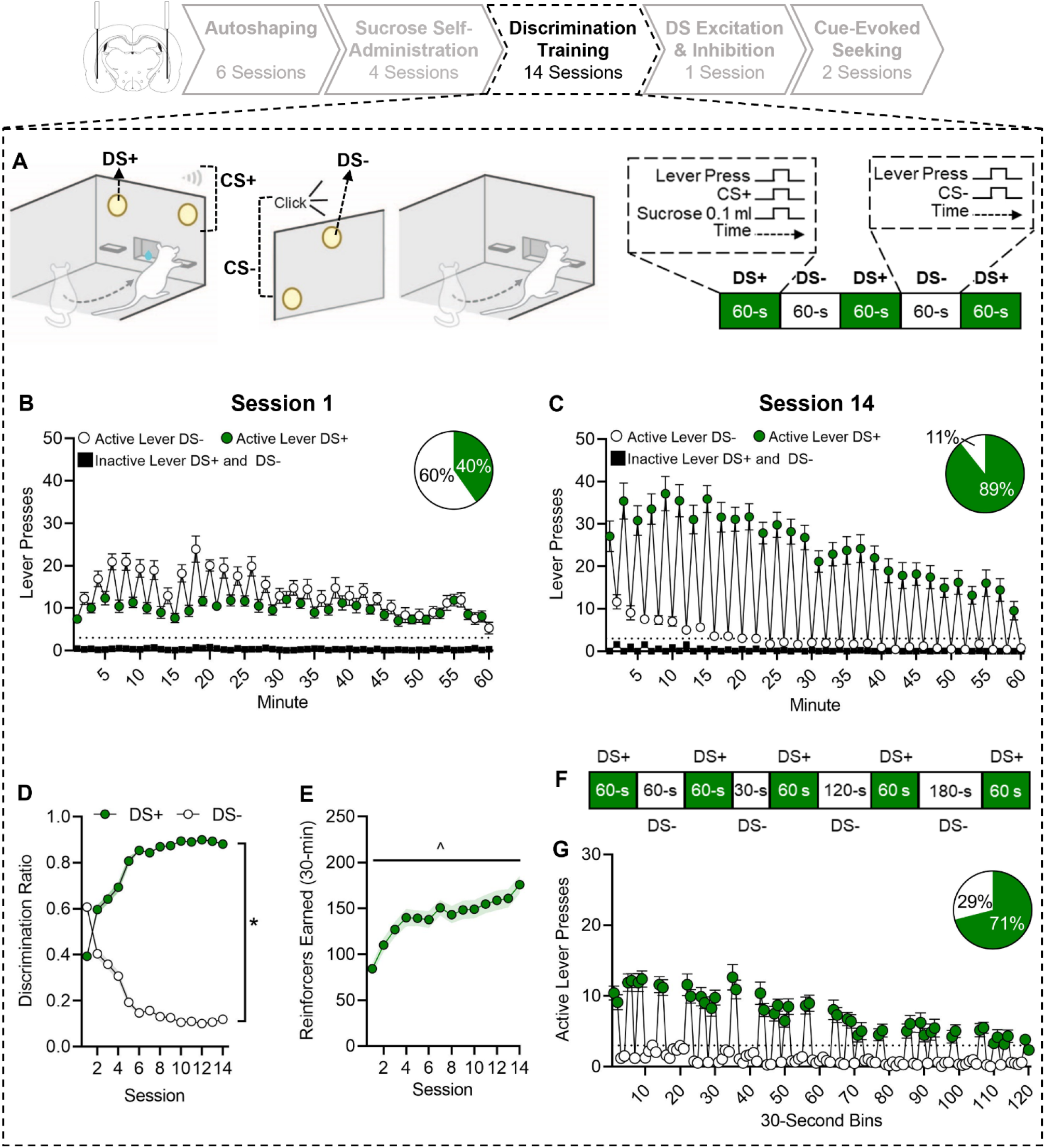
Sucrose self-administration comes under the control of discriminative stimuli. **(A)** Schematic of the discrimination procedure, adapted from Servonnet et al., 2023, with permission from Elsevier. Average (± SEM) inactive lever presses (black squares) and active lever presses made during DS+ (green circles) and DS- (white circles) presentations on **(B)** Session 1 and **(C)** Session 14. Insets represent the percentage of active lever presses made during the DS+ (green) and DS- (white). Average (± SEM) **(D)** DS+ and DS- discrimination ratios and **(E)** reinforcers earned. **(F)** Schematic of the discrimination procedure with varying DS- durations. **(G)** Average (± SEM) active lever presses made during DS+ (green circles) and DS- (white circles). The inset represents the percentage of active lever presses made during DS+ (green) and DS- (white) presentations. DS+, discriminative stimulus signaling sucrose availability. DS-, discriminative stimulus signaling sucrose unavailability. *DS+ > DS- discrimination ratio on session 14. ^ Session1 < Session 14 reinforcers earned. *n* = 16 males, 17 females.

A discrimination ratio was calculated to determine the ratio of active lever presses during DS+ versus DS-presentations [DS+ ratio: (total DS+ active lever presses/total DS+ and DS-active lever presses); DS-ratio: (total DS-active lever presses/total DS+ and DS-active lever presses)].

### Discrimination with Unpredictable DS- Duration

In the preceding sessions, DS+ and DS-presentations alternated every 1-min. As such, responding could be due to rats keeping time rather than assigning conditioned properties to the cues. To test this possibility, rats received one discrimination session (1-h) during which the DS-duration varied and was therefore unpredictable to the rats (30, 60, 120, 180-s; see Figure 3F). If responding was a function of rats tracking time, then active lever pressing should escalate after 60 s of DS-presentation. Alternatively, if the DSs controlled sucrose self-administration behaviour, then rats should inhibit lever pressing during the DS-regardless of its duration and increase their responding only upon DS+ presentation.

### Conditioned excitation by the DS+ and conditioned inhibition by the DS-

To confirm the conditioned excitation and inhibition effects of the DS+ and DS-, respectively, we determined whether the CS+ acquired conditioned reinforcing effects and whether DS+ presentation enhances these effects while DS-presentation suppresses these effects. Thus, after discrimination training, rats were divided into two groups and given a test (20-min duration) where they could lever press to obtain presentations of the CS+ alone, without sucrose (see Figure 4A). For both groups, the test session began with the active and inactive levers extending. In one group, pressing the active lever produced the CS+ on an FR3 schedule without any DS. In the second group, lever extension was followed by 2-min presentations of the DS+ and DS- in an alternating manner, during which active lever pressing produced presentations of the CS+ on an FR3 schedule. If the DSs modulate the conditioned properties of the CS+, then relative to the CS+ alone, DS-presentations should reduce responding for the CS+ and conversely, DS+ presentations should increase responding.

**Figure 4.**
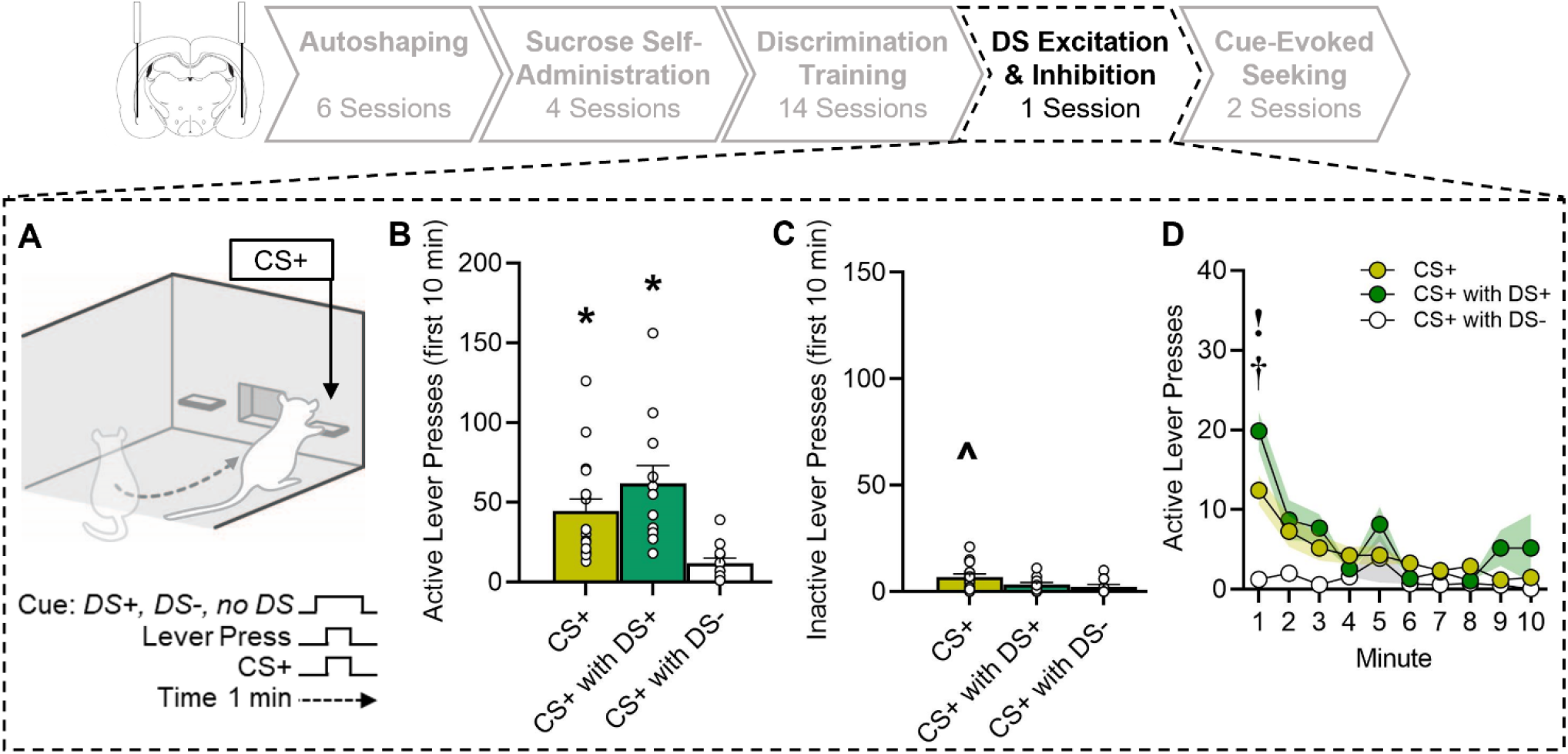
Effects of discriminative stimuli on the conditioned reinforcing properties of the conditioned stimulus. **(A)** Schematic of the conditioned reinforcement test procedure, adapted from Servonnet et al., 2023, with permission from Elsevier. Average (± SEM) **(B)** active lever presses and **(C)** inactive lever presses made to obtain CS+ presentations in the absence of the DSs (yellow) or in the presence of the DS+ (green) or DS- (white) during the first 10-min of the session. **(D)** Average (± SEM) active lever presses made across the first 10 min of the test. DS+, discriminative stimulus previously signaling sucrose availability. DS-, discriminative stimulus previously signaling sucrose unavailability. CS+, conditioned stimulus previously paired with sucrose delivery. *Active lever presses > Inactive lever presses; CS+ and CS+ with DS+ > CS+ with DS-. ^CS+ > CS+ with DS+. †CS+ and CS+ with DS+ > CS+ with DS-. ! CS+ with DS+ > CS+. n = 16 males, 17 females.

### The Effect of Intra-BLA LY379268 on Cue-evoked Sucrose Seeking

Two cue-evoked sucrose seeking tests (31-min/session) were conducted to determine the extent to which response-independent presentations of the cues trigger increases in sucrose seeking in the absence of sucrose, and the contributions of BLA mGlu_2/3_ receptor activity to this effect (Figure 5A). Sessions began with a 1-min no-cue period during which the active and inactive levers were inserted. The first cue (DS+, DS-, CS+, CS-, or DS+CS+; counterbalanced across rats) was then presented for 1-min independent of the rats’ responding, followed by a 1-min inter-trial interval (ITI) during which levers remained extended but the cue was no longer presented. This cycle continued until each cue type was presented three times in a pseudo-random order. Throughout these test sessions, lever pressing produced neither sucrose nor sucrose-associated cues. LY379268 (3 µg/hemisphere) or aCSF was bilaterally microinfused into the BLA 5-min before each test (counterbalanced). Rats received two additional discrimination training sessions between test sessions.

**Figure 5.**
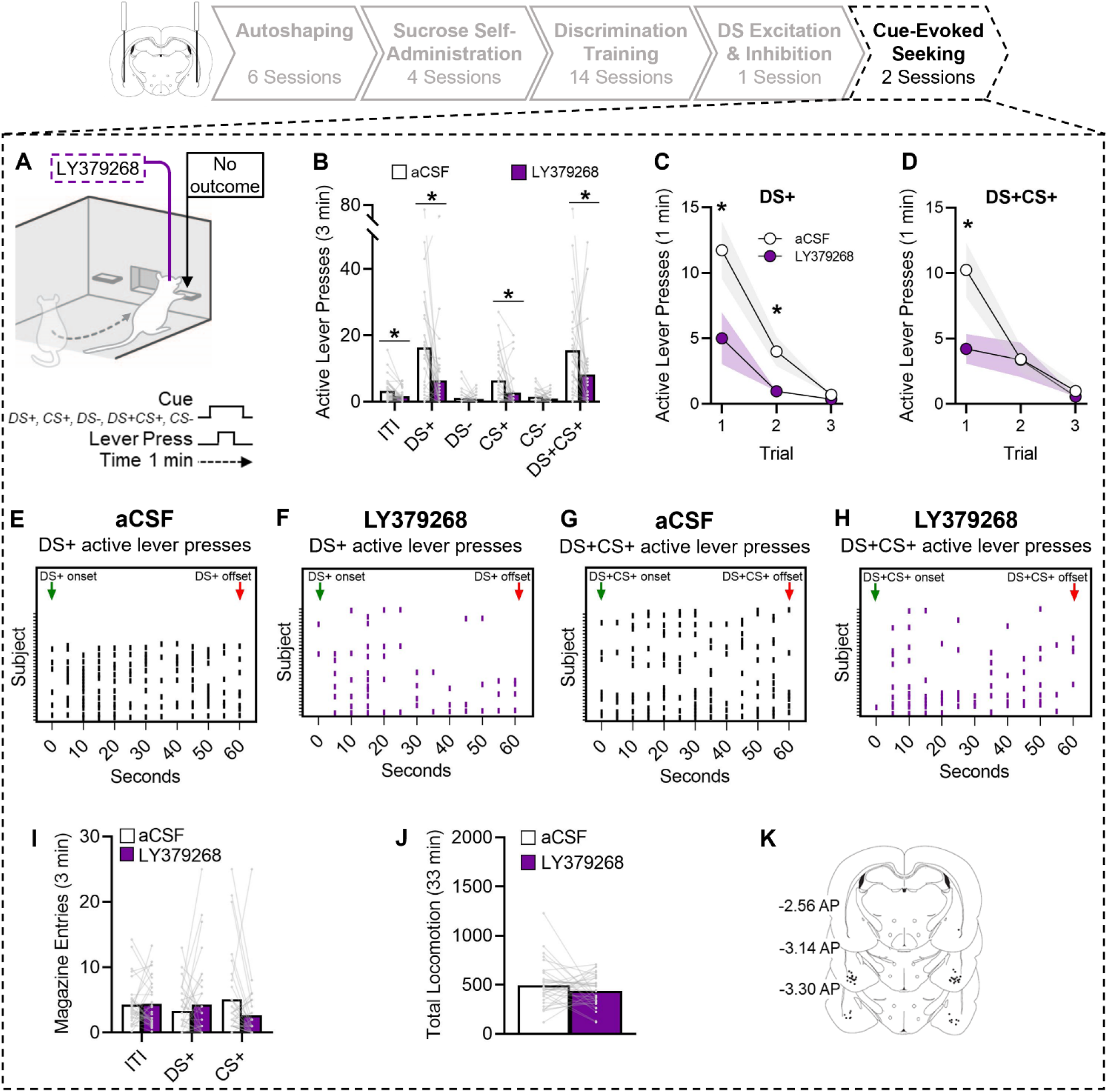
Injections of LY379268 into the basolateral amygdala (BLA) suppress the ability of a discriminative stimulus to increase sucrose seeking. **(A)** Schematic of the cue-evoked sucrose seeking test procedure, adapted from Servonnet et al., 2023, with permission from Elsevier. **(B)** Average (± SEM) active lever presses summed across the test session under aCSF (white bars) and LY379268 (purple bars) treatment. Average (± SEM) active lever presses across trials during **(C)** DS+ and **(D)** DS+CS+ presentations under aCSF (white circles) and BLA LY379268 (purple circles) treatment. Timestamps of individual active lever presses during presentation of the **(E & F)** DS+, and **(G & H)** DS+CS+ combined, under aCSF (black lines) or LY379268 treatment (purple lines). Average (± SEM) **(I)** magazine entries and **(J)** locomotion summed across the test session following aCSF (white bars) and LY379268 (purple bars) treatment. **(K)** Representation of estimated injector tip placements in the BLA. Numbers indicate anteroposterior coordinates from bregma. * aCSF > LY379268. n = 16 males, 17 females.

### Statistical Analyses

All experiments used a mixed design. The within-subject factors were treatment condition (LY379268, aCSF) and cue condition (ITI, DS+, CS+, DS-, CS-, DS+CS+). The between-subject factor was biological sex. Non-parametric tests were used because Shapiro-Wilk tests confirmed that all data sets were non-normally distributed. We conducted correlations using Spearman’s correlation tests. Based on prior work [9,15,16,19,23,51,52], we predicted that compared to the DS- and the CS+, the DS+ would trigger greater increases in lever pressing during cue-evoked seeking tests. We also predicted that responding to the DS+ alone and to the DS+CS+ combined would be similar [15]. Therefore, a priori post-hoc analyses were conducted on these comparisons using Wilcoxon tests to analyze within-subject factors, and Mann-Whitney tests to analyze between-subject factors. We used the Holm-Bonferroni correction for multiple comparisons. Statistical analyses were conducted with IBM SPSS (Version 26) using an α level of *p <* 0.05. Graphs were created with Graphpad Prism (Version 8; La Jolla, CA).

## Results

As detailed in the Supplementary text, there were no sex effects during behavioural training or testing and therefore data were collapsed across sex for analyses.

### Autoshaping

Rats increased their CS lever activations (Figure 1B; Figure 1 vs. 6: *z =* -4.53, *p <* 0.001) and kept their rate of CS magazine entries stable (Figure 1C; Figure 1 vs. 6: *p >* 0.05) during autoshaping sessions. Figure 1D depicts the conditioned response bias in sign-trackers, goal-trackers, and intermediates across autoshaping sessions. By the end of autoshaping, nearly half of the rats were classified as sign trackers, followed by intermediates, then goal trackers (Figure 1E).

### Sucrose Self-Administration

Rats pressed more on the active vs. inactive lever during sucrose self-administration sessions (Figure 2B; Session 1: *z =* -5.02, *p <* 0.001; Session 2: *z =* -5.04, *p <* 0.001; Session 3: *z =* -5.02, *p <* 0.001; Session 4: *z =* -5.02, *p <* 0.001). As Figure 2B shows, the number of active lever presses (Session 1 vs 2 and Session 3 vs 4; All p’s > 0.05), reinforcers earned (Session 1 vs 2 and Session 3 vs 4; All p’s > 0.05), and magazine entries (Figure 2C: Session 1 vs 4 *p >* .05), remained elevated across self-administration sessions and stable under each schedule of reinforcement, whereas inactive lever presses decreased across sessions (Session 1 vs 4 *z =* -3.43, *p <* 0.001). Thus, the rats rapidly learned to self-administer and retrieve sucrose from the magazine.

### Sucrose Self-Administration Under the Control of Discriminative Stimuli

On Session 1 of discrimination training, active lever presses peaked during the DS-, particularly in the first half of the session (Figure 3B). Indeed, on Session 1, rats made 60% of their active lever presses during DS-presentations and 40% during the DS+ (Figure 3B inset). However, by Session 14, responding peaked during DS+ presentations (Figure 3C), such that rats now made most of their active lever presses during the DS+ (89%; Figure 3C inset) and suppressed responding during the DS- (11%).

Figure 3D further highlights the acquisition of discriminated responding, where the DS+ discrimination ratio increased from Session 1 to 14 (*z =* -4.89, *p <* 0.001) and the DS-discrimination ratio decreased from Session 1 to 14 (*z =* -5.01, *p <* 0.001). On Session 1, discrimination ratios were higher for the DS- vs. DS+ (*z =* -4.96, *p <* 0.001). However, by Session 14, this pattern reversed, with rats showing greater discrimination ratios during the DS+ vs. DS- (*z =* -5.01, *p <* 0.001). Accordingly, Figure 3E shows that sucrose intake also increased from Sessions 1 to 14 (*z =* -5.01, *p <* 0.001). Thus, both females and males learned to respond for sucrose during DS+ presentation and to inhibit responding during the DS-.

Active lever presses may have peaked during the DS+ because the DSs controlled sucrose self-administration behaviour or, alternatively, because the rats were tracking time during the sessions, as the duration of each DS presentation (1 min) was predictable to the rats. To address this, rats received a discrimination training session during which the duration of DS-presentations varied between 30-180 s (Figure 3F). Despite the now unpredictable DS-duration, rats continued to press more on the active lever during DS+ presentation and to inhibit responding during DS-presentation, regardless of duration (Figure 3G). Thus, the DSs came to control sucrose self-administration.

### Conditioned excitation by the DS+ and conditioned inhibition by the DS-

During a conditioned reinforcement test where rats could lever press for the CS+ in the absence of sucrose, rats pressed more on the active vs. inactive lever (Figures 4B-C) when the CS+ was presented alone (*z =* -3.62, *p <* 0.001) or in the presence of the DS+ (*z =* -2.93, *p =* 0.006). However, when the DS- was present, active lever pressing for the CS+ dropped significantly (Fig. 4B; vs. CS+ alone: *z =* -3.11, *p =* 0.002; vs. with DS+ present: *z =* -2.93, *p =* 0.006), such that there was also no longer significant discrimination between active and inactive levers (*p >* 0.05). Overall rates of active lever pressing were similar when the CS+ was presented alone or with the DS+ present (*p >* 0.05). Rats also pressed more on the inactive lever when the CS+ was presented alone than when the DS+ was also present (Figure 4C; *z =* -2.60, *p =* 0.009). Thus, the DS-significantly inhibited the conditioned reinforcing effects of the CS+.

Figure 4D shows minute-by-minute responding across the first 10 min of test. During the 1st minute, the presence of the DS-suppressed responding compared to the other conditions (vs. CS+ alone: *z =* -2.98, *p =* 0.003; vs. CS+ with DS+: *z =* -3.07, *p =* 0.003), whereas rats responded more for the CS+ in the presence of the DS+ than they did for the CS+ alone (*z =* -1.96, *p =* 0.03). Figure 4D also shows that responding for the CS+ extinguished across trials, both when the CS+ was presented alone or with the DS+.

In summary, a CS paired with sucrose delivery acquired conditioned reinforcing value, enabling it to maintain responding for sucrose when the reinforcer is not available. The presence of a DS+ only increased this behaviour transiently. In contrast, the presence of a DS- significantly inhibited the conditioned reinforcing effects of the CS+ across the test session.

We found no statistically significant correlations between Pavlovian conditioned response bias (i.e., a sign- or goal-tracking phenotype) and active lever pressing during conditioned reinforcement tests (see Table 2). Thus, under our test conditions, Pavlovian conditioned responding phenotype did not significantly predict the response to CSs presented alone or in the presence of DSs.

**Table 2.** Correlations between Pavlovian conditioned response bias and behavioural outcomes.

|  | *r* | *p* |
| --- | --- | --- |
| **Conditioned reinforcement** |  |  |
| Active lever presses for CS+ | -0.31 | > .05 |
| Active lever presses for DS+/CS+ | -0.05 | > .05 |
| Active lever presses for DS-/CS+ | 0.07 | > .05 |
| **Cue-evoked sucrose seeking (aCSF)** |  |  |
| Active lever presses during DS+ | 0.26 | > .05 |
| Active lever presses during CS+ | -0.23 | > .05 |
| Active lever presses during DS- | -0.10 | > .05 |
| Active lever presses during DS+CS+ | -0.12 | > .05 |
| Active lever presses during CS- | 0.06 | > .05 |

### Cue-Evoked Sucrose Seeking

During a test where sucrose was unavailable, we assessed sucrose-seeking behaviour in the presence of the DS+, DS-, CS+, CS-, and DS+CS+. We also assessed the effects of intra-BLA injections of LY379268 on this behaviour. Figure 5K shows estimated injector tip placements in the BLA.

### Cue-evoked sucrose seeking following intra-BLA aCSF injections

Relative to the inter-trial interval (ITI), rats responded significantly more on the active lever during presentation of the DS+ (*z =* -4.19, *p <* 0.001; Figure 5B) and DS+CS+ combined (*z =* -4.49, *p <* 0.001; Figure 5B). Relative to the ITI, CS+ presentation did not change rates of active lever pressing (*p >* 0.05) and both the DS- (*z =* -3.49, *p <* 0.001) and CS- (*z =* -2.65, *p =* 0.02) reduced responding. There were also differences in responding between cue types. Compared to the CS+, the DS+ presented alone (z =-3.19, *p =* 0.001) or in combination with the CS+ (*z =* -3.87, *p <* 0.001) triggered significantly more sucrose-seeking behaviour. Thus, relative to baseline, presentations of the DS+ and DS+CS+ evoked increases in sucrose seeking, the CS+ had no effect, and the DS- and CS- both suppressed sucrose seeking.

There were no significant correlations between Pavlovian conditioned response bias and sucrose seeking behaviour triggered by any cue type (see Table 2). Thus, a sign- vs. goal-tracker phenotype did not predict the ability of reward-associated CSs or DSs to evoke sucrose-seeking behaviour.

### Cue-evoked reward seeking following intra-BLA LY379268 injections

As Figure 5B shows, compared to aCSF, intra-BLA LY379268 significantly reduced the rate of active lever pressing during the ITI (*z =* -2.64, *p =* 0.01) and in response to the DS+ (*z =* -3.17, *p =* 0.01), CS+ (*z =* -2.96, *p =* 0.01), and DS+CS+ combined (*z =* -2.79, *p =* 0.02). There were no treatment effects on responding to the DS- or CS- (all p’s > 0.05). Thus, activation of mGlu_2/3_ receptors in the BLA reduced sucrose-seeking behaviour during the ITI and the CS+, and it also suppressed the ability of DS+ and DS+CS+ presentation to boost sucrose-seeking responses.

To further examine the ability of intra-BLA LY379268 to suppress increases in sucrose seeking triggered by the DS+, we analysed trial-by-trial responding (Figures 5C-D; each cue type was presented 3x/test session). Compared to aCSF, LY379268 significantly reduced the increases in sucrose seeking evoked by the DS+ during the first and second trial (Trial 1; *z =* -2.57, *p =* 0.02; Trial 2 DS+: *z =* - 2.89, *p =* 0.01), and the DS+CS+ during the first trial (Trial 1; *z =* -2.68, *p =* 0.007). Thus, trial-by-trial analysis confirms that activation of BLA mGlu_2/3_ receptors suppresses the increased sucrose seeking triggered by a DS+, whether the DS+ is presented alone, or with the CS+.

Individual data further highlight the effects of intra-BLA LY379268 administration. Figures 5E & G illustrate responding to the first DS+ and DS+CS+ trials following aCSF treatment in individual rats. Rats typically showed sustained increases in sucrose-seeking behaviour during DS+ and DS+CS+ presentation. In these same rats, LY379268 treatment significantly suppressed responding (Figures F & H).

Finally, compared to intra-BLA aCSF, LY379268 did not significantly change magazine entries (Figure 5I; *p >* 0.05) or locomotion (Figure 5J; *p >* 0.05) during the test session. Thus, activation of BLA mGlu_2/3_ receptors produced no significant motor effects [45,53].

## Discussion

Using a new method to assess how DSs and CSs influence reward-seeking behaviour, we found that mGlu_2/3_ receptor activity in the BLA suppresses the increased reward pursuit otherwise produced by a DS+. Rats first learned to self-administer sucrose during DS+ presentations and to suppress responding during DS-presentations within the same session. Self-administered sucrose was also paired with a CS+, which gained reinforcing value and maintained sucrose-seeking behaviour in the absence of sucrose reward. DS-presentation inhibited this conditioned reinforcing effect, further confirming its inhibitory properties. When sucrose was unavailable, presentation of the DS+ alone or with the CS+ increased sucrose seeking, while the CS+ alone was ineffective. Finally, infusions of the mGlu_2/3_ receptor agonist LY379268 into the BLA suppressed the increase in sucrose seeking triggered by the DS+.

### A new procedure to characterize DS+ and CS+ effects on reward seeking

Rats learned to respond for sucrose during trials where a DS+ signaled sucrose availability and to inhibit responding during trials where a DS- signaled sucrose unavailability. During each self-administration session, we presented the DS+ and DS- in discrete, 1-min intervals, in a predictable pattern. As such, the peaks in responding during DS+ trials and the troughs in responding during DS- trials could be because the DSs came to control sucrose self-administration or, alternatively, because rats were keeping time. However, the rats continued to load up on sucrose during the DS+ and supress responding during the DS- even when we varied the duration of DS- presentation, making it unpredictable to the rats (see also [15]). Thus, our new procedure allows DSs to gain conditioned excitatory (DS+) and inhibitory (DS-) properties, such that the DSs come to control the pursuit of sucrose. This procedure adds to a recently developed trial-based procedure to assess DS controlled drug or food self-administration [20,21], with the added benefit of enabling direct comparison of DS vs. CS effects on reward seeking in the same subjects within the same session.

After training, presentation of the DS+ and DS+CS+ combined triggered increases in sucrose seeking relative to baseline, while the CS+ did not change responding relative to baseline. These findings are consistent with others showing that via associative pairings with reward availability, response-independent presentation of a DS+ robustly increases reward-seeking actions [15,16,20–23], whereas a CS+ is ineffective [9,15–17,19], or minimally effective [18]. The DS+ may have induced a conditioned craving state that stimulated sucrose-seeking responses [54]. Alternatively, the DS+ might have triggered a cognitive expectation of sucrose, leading to increased motivation or craving for sucrose reward. In agreement, rates of craving are at their peak when subjects expect that the coveted reward will soon be available [55].

Our new procedure addresses important caveats of previous research. Previous studies have often compared DS vs. CS effects in separate test sessions or separate rats [9,51]. By presenting all cue types to all rats during a single test session, our procedure mitigates potential re-resting effects including extinction of the lever pressing response and/or of cue-reward associations. Our procedure also enables the identification of common as well as unique effects of DSs and CSs on brain and behaviour, in the same subjects within the same test session. While some studies have presented DSs and CSs in the same test session [16], they were presented during long, 30-minute intervals. In our procedure, both DSs and CSs are presented in discrete, short trials. This facilitates the application of event-locked measurements and manipulations of neuronal activity. Our procedure also allows us to identify and account for individual variability in the response to a DS or CS. Finally, we have validated our procedure across the sexes, demonstrating effective discriminated self-administration and DS-controlled reward seeking in both females and males. Thus, our approach provides a valuable tool for future research on the neurobiological mechanisms underlying DS and CS effects on reward seeking.

### Conditioned excitation and inhibition of the reinforcing properties of a CS+

Rats pressed a lever, previously associated with sucrose delivery, to earn presentations of a CS+, thus maintaining reward-seeking actions through conditioned reinforcement [9–15]. DS+ presentation invigorated the pursuit of the CS+, confirming its role as a conditioned excitor. However, consistent with prior work [56,57] the effect was short-lived, extinguishing after the first minute. In parallel, DS- presentation suppressed lever pressing for the CS+ quite persistently, also confirming its role as a conditioned inhibitor. Thus, the DS- inhibited the conditioned motivational salience of a reward-paired CS+, reducing its ability to maintain instrumental pursuit [54]. Prior work shows that a DS- can also reduce cocaine seeking evoked by a DS+ [15,20,58,59]. Our results extend this literature by demonstrating that the conditioned inhibitory effects of a DS- extend to reward seeking maintained by a CS+. This could have clinical applications, as cues signalling reward unavailability could potentially be used as conditioned inhibitors of pathological reward-seeking behaviour in eating, gambling, and substance use disorders [58,60,61].

### Sign- versus goal-trackers

Others have observed that a sign-tracking phenotype predicts increased CS+ guided reward seeking [62] while a goal-tracking phenotype predicts increased DS+ guided reward seeking [47,48]. In contrast, and similar to others [63–65], we found that Pavlovian conditioned phenotype did not significantly predict responding to the DS+, CS+ or DS+CS+. Methodological parameters could explain the differences between studies. Prior work used extinction training before reward-seeking tests [62], whereas we did not. Prior studies also tested CS vs. DS effects separately [48,63], whereas we did so in the same subjects, within the same test sessions.

### Activating BLA mGlu_2/3_ receptors reduces DS+ evoked sucrose seeking

Intra-BLA administration of LY379268 suppressed DS+ evoked increases in sucrose seeking, without producing overt motor effects [45,53]. Intra-BLA infusions of LY379268 could be acting by attenuating the predictive properties of the DS+. This is unlikely as LY379268 did not influence magazine entries elicited by the DS+ (or CS+), and cue-triggered magazine entries are thought to reflect cue predictive value [66,67]. Alternatively, mGlu_2/3_ receptor activity in the BLA might attenuate the motivational salience of the DS+, thus reducing its ability to trigger wanting of sucrose reward [54]. This is consistent with findings that intra-BLA infusions of LY379268 reduce the ability of a CS+ to invigorate reward seeking in a Pavlovian-to-instrumental-transfer (PIT) task [45]. PIT is similar to our cue-evoked seeking task as both involve response-independent cue presentation, thus measuring the ability of reward-associated cues to trigger a Pavlovian conditioned motivational state of wanting for the associated reward.

The BLA is a critical hub for the ability of reward cues to attract attention, trigger, reinforce and invigorate reward seeking, and glutamatergic inputs from the cortex and thalamus mediate these cue effects [43,44]. When activated, mGlu_2/3_ receptors reduce pre-synaptic glutamate release to attenuate glutamate neurotransmission [38–40,68,69]. LY379268 is reported to have no effects of its own on extracellular glutamate levels but to suppress pharmacologically-evoked glutamate, at least in the medial prefrontal cortex [70]. To the extent that these findings translate to the BLA, it would suggest that activation of BLA mGlu_2/3_ receptors suppresses the increase in glutamate neurotransmission triggered by the reward-associated DS+. In agreement, using a PIT procedure, Malvaez et al [43] found that reward-predictive cues elevate glutamate concentrations in the BLA thus contributing to cue-triggered increases in reward-seeking actions. Thus, we propose that intra-BLA infusion of LY379268 suppresses the increases in reward seeking produced by a DS+ by reducing glutamate release and neurotransmission in the BLA and in downstream BLA-dependent circuits. These may include BLA projections to the nucleus accumbens core, medial prefrontal cortex, and ventral hippocampus [27,30,31,71–76].

## Conclusions

We developed a behavioural assay to identify common and unique effects of reward associated DSs and CSs in the same rats, within the same test session. Using this procedure, we found that compared to a CS indicating reward delivery, a DS signalling reward availability is more effective in triggering increases in reward-seeking responses, illustrating the power of DSs over appetitive behaviour (see also [15,16,20–23]). We also found that activity at mGlu_2/3_ receptors in the BLA supresses the increases in reward-seeking behaviour triggered by a DS+ when sucrose reward is unavailable. We conclude that glutamatergic neurotransmission within the BLA mediates the ability of reward-predictive DSs to enhance reward-seeking actions.

## Acknowledgements

This research was funded by grants to A.-N.S. from the National Science and Engineering Research Council of Canada (Grant 355923) and the Courtois Fund. M. R. L. is supported by a postdoctoral fellowship from the Natural Sciences and Engineering Research Council of Canada. A.M.B. is supported by a Merit scholarship from the Université de Montréal.

