## Supplemental Materials for "Activation of Group II Metabotropic Glutamate Receptors in the Basolateral Amygdala Inhibits Reward Seeking Triggered by Discriminative Stimuli"

**Apparatus**

Behavioural procedures were conducted in conditioning chambers with standard grid floors (Med Associates Inc., St-Albans, VT) enclosed in sound-attenuating, ventilated melamine cubicles (ENV-016MD). The chamber back wall held a sound generator (top right; ENV-223AM; 2900 Hz, 75 dB), and two white cue lights (one top center; ENV-215M, one bottom left; SciFab, Ann Arbor, MI, USA). On the front wall, two retractable levers (ENV-112CM) were located on either side of a magazine (ENV-200R3M), into which 0.1 ml of a 10% *w*/*v* liquid sucrose solution (CAS#57501, Bioshop; sucrose dissolved in tap water) was delivered via a liquid dispenser (ENV-201A) connected to a 20-ml syringe. Entries into the magazine were measured via interruptions of two infrared motion detectors located respectively, at the bottom and top of the magazine opening. During sucrose self-administration and discrimination training, presses on the active lever (counterbalanced randomly between the left- and right-side levers) delivered liquid sucrose. A white cue light (ENV-221M) was located above each lever. A clicker sound generator (made in-house) was located outside of the chamber. Horizontal locomotor activity within the chambers was measured via interruptions of four infrared beam sensors mounted 2.5-cm above the grid floor. Med PC IV software on a PC delivered stimuli and recorded behavioural responses.

**Stereotaxic Surgery**

Rats underwent stereotaxic surgery using standard procedures to bilaterally implant stainless steel cannulae (26 gauge; HRS Scientific) into the basolateral amygdala (BLA; -2.8 mm anterior-posterior, ± 5.0 mm medial-lateral relative from Bregma, and -6.4 mm ventral from the skull surface (Garceau et al., 2021; Servonnet et al., 2020). During intracranial drug microinfusions, the injector tip (33 gauge, HRS Scientific) protruded 2.0 mm below the cannula base, resulting in a final ventral coordinate of -8.4 mm. Guide cannulae were occluded with dummy cannulae attached to a dustcap. Rats received perioperative carprofen (15 mg/kg, subcutaneous; CDMV, QC, Canada) and penicillin (60 mg/kg, subcutaneous; Derapen, CDMV, QC, Canada) to prevent pain and infection, respectively. Behavioural training began after a 1-week recovery period.

**Sucrose Habituation**

Following bilateral implantation of cannulae into the BLA and 1 week of recovery from stereotaxic surgery, rats were habituated to drinking the sucrose solution by receiving 48-h access to a water bottle and a separate bottle containing 80 ml of 10% *w/v* sucrose in the home cage.

**Water Restriction**

The day following sucrose habituation, water restriction began to promote sucrose self-administration behaviour. Rats had 6 hours/day of water access for the first 4 days, then 4 hours/day for 3 days, then 2 hours/day until the end of the experiment. Rats received access to water at least 1 h after behavioural training.

**Magazine Training**

Two days after sucrose habituation, rats received two magazine training sessions (30-min duration) to become familiar with receiving sucrose in the magazine. Sessions began with the fan turning on, followed by delivery of 0.1 ml of sucrose solution on a VI 45-s schedule (20 - 70-s intervals). A total of 30 sucrose deliveries were given per session. The magazines were checked at the end of all behavioural training sessions to verify that the sucrose solution was retrieved. By the end of the second magazine training session, and throughout subsequent training phases, all rats drank the sucrose.

**Intra-Basolateral Amygdala LY379268** **Microinfusions**

LY379268 solutions were prepared by dissolving LY379268 (Tocris, CAT#2453) in artificial cerebral spinal fluid (aCSF) to obtain a 3.0 µg/0.5 µl concentration. The solution was solubilized via sonication at 45ºC for 10 min. Bilateral microinfusions were administered into the BLA with a 33-gauge injector attached to polyethylene tubing connected to a 5-µL Hamilton syringe (VWR, 88000). Microinfusions were delivered by a syringe pump (Harvard Apparatus, PHD 2000) at a rate of 0.3 µl/min and injectors remained in place for 1-min to promote drug diffusion.

**Histology**

At the end of all behavioural tests, rats were transcardially perfused with paraformaldehyde and saline and coronal sections (40 µm) were collected from paraformaldehyde-fixed brains using a cryostat (−20°C) for light microscopy using a standard protocol (Chaudhri et al., 2013). Ventral placements of injector tips were identified using the Paxinos and Watson rat brain atlas (Watson & Paxinos, 2007). One female rat was removed from statistical analyses due to cannulae misplacement.

**Females and Males Showed Similar Pavlovian Conditioned Phenotype**

There were no sex differences in the pattern of responding across autoshaping sessions (Data not shown; Session 1; CS lever activations: *z* = -0.36, *p* = 0.74; CS magazine entries: *z* = -0.09, *p* = 0.93; Session 6; CS lever activations: *z* = -0.22, *p* = 0.85; CS magazine entries: *z* = -1.03, *p* = 0.31). The distribution of sign-trackers, goal-trackers, and intermediates was similar in females and males (see figure below).

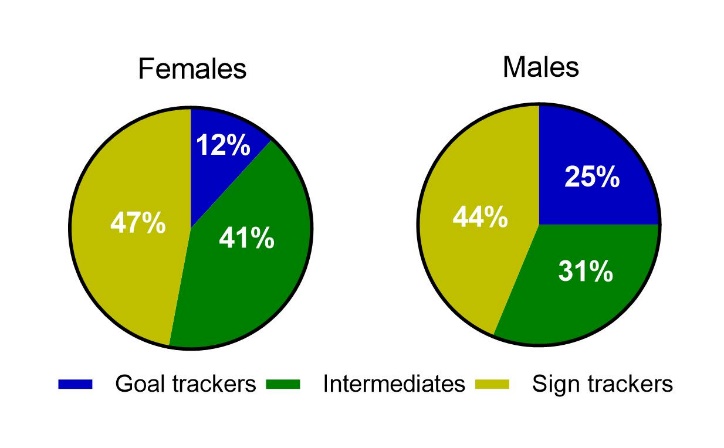

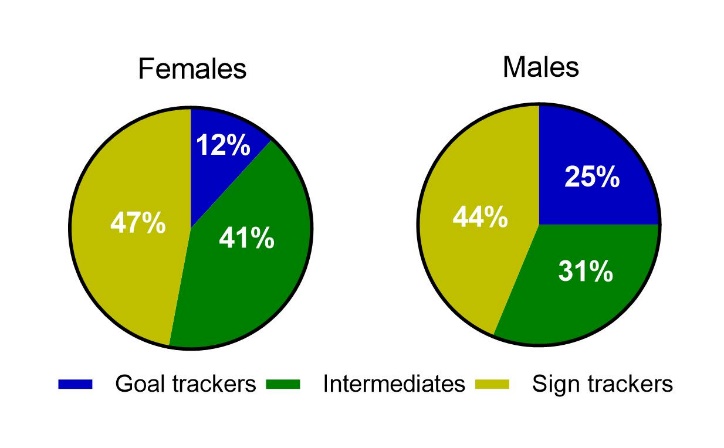

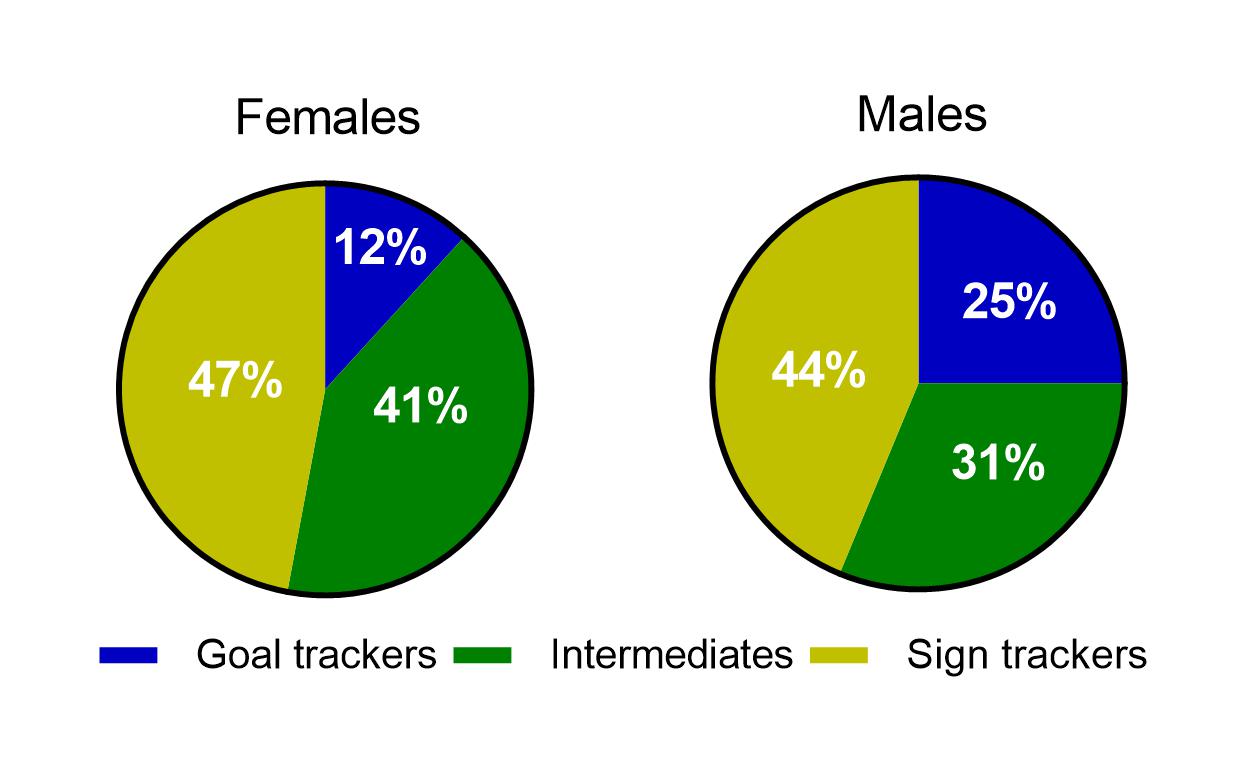

**Females and Males Showed Similar Sucrose Self-Administration**

Females and males pressed for sucrose at similar rates (data not shown; Session 1: *z* = 2.03, *p* = 0.26; Session 2: *z* = 2.03, *p* = 0.26; Session 3: *z* = 1.73, *p* = 0.40; Session 4: *z* = 2.03, *p* = 0.26). Females and males earned a similar number of reinforcers, save for Session 2 where males earned more (data not shown; Session 1: *z* = 0.27, *p* = 0.79; Session 2: *z* = 2.60, *p* = 0.04; Session 3: *z* = 0.58, *p* = 0.74; Session 4: *z* = 1.34, *p* = 0.35). Finally, females and males entered the magazine at similar rates (data not shown; Session 1: *z* = 0.23, *p* = 0.82; Session 2: *z* = 1.50, *p* = 0.14; Session 3: *z* = 1.19, *p* = 0.25; Session 4: *z* = 0.33, *p* = 0.77).

**Females and Males Showed Similar Sucrose Self-Administration under Discrimination Stimuli**

There were no sex differences in the rates of active lever pressing during DS+ and DS- presentation (Data not shown; Sessions 1: DS+: *z* = 1.95, *p* = 0.05; DS-: *z* = 2.34, *p* = 0.08; Session 14: DS+: *z* = 2.27, *p* > 0.05; DS-: *z* = 1.46, *p* = 0.15), or in discrimination ratios (data not shown; Session 1: DS+: *z* = 0.83, *p* = 0.42; DS-: *z* = -0.83, *p* = 0.42; Session 14: DS+: *z* = 1.00, *p* = 0.33; DS-: *z* = -1.00, *p* = 0.33).

**Females and Males Showed Similar Responding for Presentations of the Conditioned Stimulus**

Females and males showed similar rates of active lever pressing to obtain the CS+ alone (*z* = -0.53, *p* = 0.61), to obtain the CS+ in the presence of the DS+ (*z* = -0.61, *p* = 0.630), and in the presence of the DS- (*z* = 0.92, *p* = 0.28).

**Females and Males Showed Similar Cue-induced Sucrose Seeking Behaviour**

After aCSF injections, females and males showed similar rates of active lever pressing across cue types (Data not shown; DS+: *z* = -1.48, *p* = 0.15; CS+: *z* = -1.43, *p* = 0.16; DS-: *z* = -0.55, *p* = 0.66; DS+CS+: *z* = -1.28, *p* = 0.22; CS-: *z* = 1.09, *p* = 0.33). The sexes also showed similar rates of responding after LY379268 injections (Data not shown; DS+: *z* = -1.48, *p* = 0.15; CS+: *z* = -2.87, *p* = 0.19; DS-: *z* = -0.94, *p* = 0.42; DS+CS+: *z* = -1.00, *p* = 0.33; CS-: *z* = -0.13, *p* = 0.93).
